# Macrophage Activation by a Substituted Pyrimido[5,4-*b*]Indole Increases Anti-Cancer Activity

**DOI:** 10.1101/581710

**Authors:** Joseph Hardie, Javier A. Mas-Rosario, Siyoung Ha, Erik M. Rizzo, Michelle E. Farkas

## Abstract

Immunotherapy has become a promising new approach for cancer treatment due to the immune system’s ability to remove tumors in a safe and specific manner. Many tumors express anti-inflammatory factors that deactivate the local immune response or recruit peripheral macrophages into pro-tumor roles. Because of this, effective and specific ways of activating macrophages into anti-tumor phenotypes is highly desirable for immunotherapy purposes. Here, the use of a small molecule TLR agonist as a macrophage activator for anti-cancer therapy is reported. This compound, referred to as PBI1, demonstrated unique activation characteristics and expression patterns compared to treatment with LPS, through activation of TLR4. Furthermore, PBI1 treatment resulted in anti-tumor immune behavior, enhancing macrophage phagocytic efficiency five-fold versus non-treated macrophages. Additive effects were observed via use of a complementary strategy (anti-CD47 antibody), resulting in ∼10-fold enhancement of phagocytosis, suggesting this small molecule approach could be used in conjunction with other therapeutics.

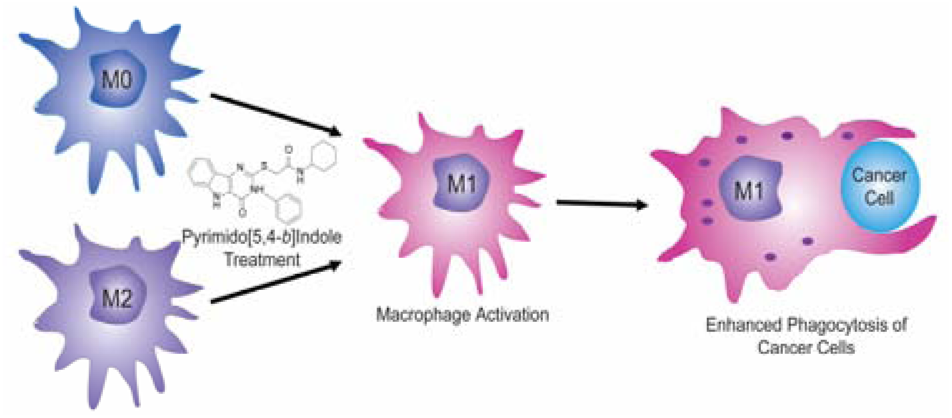

## 1. Introduction

Cancer is one of the leading causes of death around the world, and fighting it is a focus of research programs globally [1]. However, most cancer therapy regimens continue to rely on non-specific chemotherapy and radiotherapy to eliminate tumors, which have severe side effects. They are also ineffective against many cancer types and can increase disease recurrence [1]. As a result, alternative strategies are sought after, such as immunotherapy [2]. Cancer immunotherapy involves utilizing the immune system to eliminate the disease, and is attractive on account of the immune system’s specificity and biocompatibility compared to traditional cancer therapy. Several immunotherapies have been approved clinically, and others have reached the clinical trial stage [3].

While immunotherapy is alluring, developing immune-based strategies is challenging. Normally, the immune system detects and eliminates pre-oncogenic cells [4], however, cancer cells can generate cytokines and receptors for immune evasion and reprogramming [5]. In this manner, tumorigenic cells are able to escape detection and disable pro-inflammatory behaviors. An example of the former is over-expression of “don’t eat me” surface marker CD47 by cancer cells, preventing phagocytosis [6], while in the latter the tumor releases chemo-attractants and anti-inflammatory signals, such as IL-4, IL-10, CS1, CSF1R, and MFG-E8, to reprogram immune cells to perform pro-tumorigenic roles [4,7]. These include facilitation of angiogenesis, epithelial to mesenchymal transition (EMT), and microenvironment remodeling [8]. Once this point is reached, the likelihood of patient survival decreases sharply [9].

Because of the immune system’s role in cancer progression, there is great interest in the re-education of immune elements into anti-cancer entities. Polarization of macrophages into M1 (classically activated) phenotypes is important toward refocusing the immune system for eliminating cancer [10]. In this immune-stimulating phenotype, macrophages attack and phagocytose tumor cells. This facilitates a larger overall attack by the immune system, resulting in tumor elimination [10]. Macrophages also generate reactive oxygen species (ROS) and present tumor antigens, which recruit T cells and B cells to the tumor site [11]. In contrast, immune-suppressing M2 (alternatively activated) phenotypes, or tumor-associated macrophages (TAMs), contribute to tumor progression, joining the tumor mass and microenvironment [10]. TAMs release pro-tumor growth factors, such as vascular endothelial growth factor (VEGF), promote vascularization, remodel the microenvironment, and silence the immune response [12,13,14]. Overall, re-programming macrophages into M1-like states and away from M2/TAM phenotypes has great potential as an anti-cancer immunotherapeutic approach [15].

The polarization of macrophages toward M1 phenotypes is a well-studied phenomenon, with known pathways identified. Specifically, the interferon gamma (IFN-γ) receptor (IFNGR) and the toll-like receptor (TLR) class are understood to activate macrophages into pro-inflammatory roles [16]. The most common strategies for *in vitro* M1 macrophage polarization involve treatment with IFN-γ (ligand for IFNGR) and lipopolysaccharide (LPS, a TLR4 agonist originating from bacterial cell walls) [16]. While these agonists are useful as research tools, both IFN-γ and LPS have drawbacks that make them non-viable therapeutically. IFN-γ is a small protein, difficult to consistently modify and incorporate into delivery vehicles [17], while LPS is a bacterial cell wall component consisting of a mixture of structures and may be contaminated with other bacterial components, resulting in off-target immune effects [18]. Systemic administration of these agents results in immune overstimulation, leading to negative outcomes including septic shock, cytokine storms, and death [18].

Because of these issues, there is a need to identify alternative macrophage activators. Additionally, therapeutic candidates should be amenable to chemical modifications and association with targeted delivery vehicles. To date, there are few immunemodulating compounds approved clinically. One example is the TLR7 activator imiquimod, which is approved for the topical treatment of genital warts and basal cell carcinoma [19]. However, in terms of general anti-cancer agents, systemically delivered drugs are desirable to provide access to a range of tumor locations and facilitate immune cell recruitment. There have been examples of anti-inflammatory antibody blockades utilized to reprogram macrophages for cancer therapy [7,20], but in terms of modifications, antibodies are more difficult to alter than small organic molecules, and are also linked to uncontrollable immune-based toxicity [21].

Here, we utilize a small molecule TLR4 activator to induce the M1 phenotype and enhance anti-oncogenic properties in macrophages (**Fig. 1**). This molecule, a pyrimido[5,4-*b*]indole referred to as **PBI1**, was previously identified among a series of compounds that activate TLR4 [22]. This molecule was shown to bind to TLR4 and induce expression of various pro-inflammatory cytokines in dendritic cells [22]. Preclinical studies have shown that structurally similar molecules are effective immune adjuvants for influenza therapy [23], however, these compounds have not been evaluated in terms of macrophage activation and anti-cancer activity. Our hypothesis is that this small molecule TLR4 agonist can activate and polarize macrophages into an anti-cancer phenotype nearly as well as naturally occurring cytokines or existing adjuvants. We demonstrate that **PBI1** upregulates pro-inflammatory genes in macrophages and induces M1-associated phenotypic changes and cytokine production. Macrophages treated with **PBI1** demonstrate enhanced anti-tumor activity toward B-cell lymphoma cells as determined by phagocytosis assays. We also show that treatment of M2-like macrophages with **PBI1** results in their re-education of macrophages toward an M1-like phenotype.

**Fig. 1.**
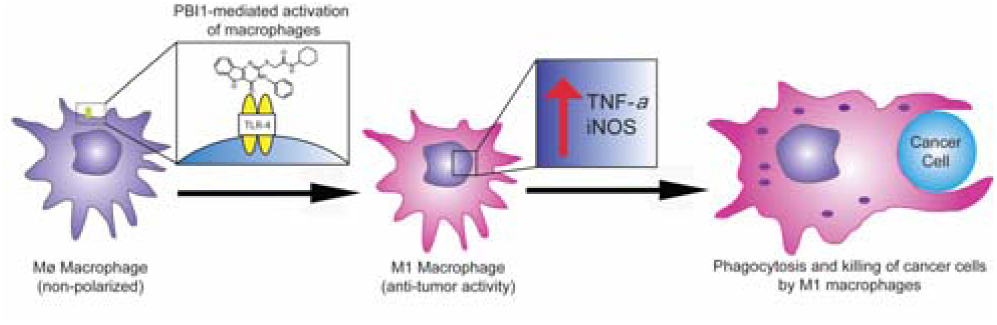
Illustration of pyrimido[5,4-*b*]indole (**PBI1**)-mediated macrophage activation. Following TLR4 stimulation, resulting M1 macrophages have enhanced levels of TNF-α and iNOS, increased ROS generation, and phagocytosis efficiency

## 2. Materials and Methods

All reagents were purchased from Thermo-Fisher Scientific except where otherwise noted. All DMSO utilized was cell culture grade (Sigma). Confocal microscopy images were obtained on an Eclipse Ti-E microscope (Nikon, Tokyo, Japan) using a 63X or 20X objective at 25 °C. Images were acquired and processed using NIS-Elements and ImageJ. Flow cytometry analysis was performed on a BD LSRFortessa 5L flow cytometer equipped with FACSDiva (BD Sciences, USA) at the Flow Cytometry Core Facility at the University of Massachusetts, Amherst. RTPCR data was generated using a CFX Connect Real-Time PCR Detection System (Biorad, Hercules, CA). For assays requiring absorbance measurements, a SpectraMax M2 plate reader was used (Molecular Devices, San Jose, CA).

### 2.1 Cell Culture

RAW 264.7 cells were purchased from American Type Culture Collection (ATCC, Manassas, VA). Primary immortalized macrophages were a gift from Prof. Susan Carpenter at the University of California, Santa Cruz. Both types of cells were cultured at 37 °C under a humidified atmosphere containing 5% CO2. Standard growth media consisted of high glucose Dulbecco’s Modified Eagle Medium (DMEM) supplemented with 10% fetal bovine serum (FBS) and 1% antibiotics (100 µg/ml penicillin and 100 µg/ml streptomycin). Daudi cells were a gift from Prof. Vincent Rotello at the University of Massachusetts Amherst and were cultured in Roswell Park Memorial Institute media (RPMI 1640) supplemented with 10% fetal bovine serum and 1% antibiotics (100 µg/ml penicillin and 100 µg/ml streptomycin). Under the above culture conditions, the cells were sub-cultured approximately once every four days and only cells between passages 7 and 15 were used for all the experiments.

### 2.2 Synthesis

The synthesis of PBI1 was largely performed as previously described [22]. A complete description of procedures and analytical data may be found in the supporting information.

### 2.3 RT-PCR Preparation

RAW 264.7 cells were plated in 24 well plates at a density of 100,000 cells/well. PBI1-treated cells were dosed with 20 µg/mL compound for 24 hours. M1 treatment group cells were treated with 50 ng/mL IFN-γ for 24 hours after which LPS was added to a final concentration of 50 ng/mL for an additional 24 hours. M2 group cells were treated with 50 ng/mL IL-4 for 48 hours. For the M2+PBI1 group, the cells were treated with 50 ng/mL IL-4 for 24 hours. 24 hours after IL-4 dosing, the cells were treated with 20 µg/mL PBI1 for 24 hours. Each experiment included three biological replicates per treatment condition. 20 µg/mL of PBI1 was chosen for this assay and several others as PBI1 demonstrated no toxicity at this concentration (Figure S10). Following treatments, RNA was extracted following the procedure below.

### 2.4 RNA Extraction and cDNA Conversion

Approximately 1.5 µg RNA was directly harvested from cells using the PureLink RNA Mini Kit (Ambion) following the manufacturer’s instructions. SuperScript IV Reverse Transcriptase, RNaseOut, 10 mM dNTPs, and 50 μM Random Hexamers were used for the conversion of approximately 150 ng of RNA to cDNA (ThermoFisher, Pittsburgh, PA), also following the manufacturer’s instructions with a sample volume of 20 µL. Briefly, primers were annealed to RNA at 65 °C for 5 minutes. Then the annealed RNA was combined with the reaction mix (containing 500 units of reverse transcriptase per sample) and amplified at 53 °C for 10 minutes and melted at 80 °C for 10 minutes. The cDNA was frozen at −20 °C and then used for RT-PCR within 1 week. RNA and cDNA were quantified using a NanoDrop 2000 (ThermoFisher). RNA and cDNA contamination, inhibition and integrity was assessed by analyzing the A260/A280 ratio, where 1.8 for DNA and 2.0 for RNA was considered “pure.”

### 2.5 Quantitative RT-PCR

RT-PCR was performed on cDNA as prepared above using a CFX Connect real-time system (Biorad) with iTaq Universal SYBR Green Supermix (Biorad, Hercules, CA). All DNA primers were purchased from Integrated DNA Technologies (Caralville, Iowa). The following primer sequences (as determined by NCBI primer-BLAST) were used: β-actin (ACTB, accession number: NM_007393, amplicon length: 86 pairs, exon 6) (forward) GATCAGCAAGCAGGAGTACGA, (reverse) AAAACGCAGCGCAGTAACAGT; iNOS (NOS2, accession number: NM_010927.4, amplicon length: 127 pairs, exon 2,3, splice variants: 1-3) (forward) GTTCTCAGCCCAACAATACAAGA, (reverse) GTGGACGGGTCGATGTCAC; TNF-α (TNF, accession number: NM_013693.3, amplicon length: 145 pairs, exon 3,4) (forward) CCTGTAGCCCACGTCGTAG, (reverse) GGGAGTCAAGGTACAACCC. 200 nM of each primer were mixed with 1 µl (100 ng) of cDNA, 10 µL SYBR green supermix and H2O to a final volume of 20 µL. Analyses were performed as follows: the samples were first activated at 50 °C for 2 min, then 95 °C for 2 min. Then denaturing occurred at 95 °C for 30 s followed by annealing at 57 °C; the denature/anneal process was repeated over 40 cycles. Relative gene expression was determined by comparing the Ct value of the gene of interest to that of the β-actin housekeeping gene, by the 2^ΔΔCt^ method [24]. Three biological replicates were performed for each treatment condition and three technical replicates were used for each biological replicate. There was no amplification for the NTC. Data was analyzed using the CFX Manager 3.1 software. CQ values were generated by using the point at which the sample fluorescence value exceeded the software’s default threshold value. Each sample was normalized to the untreated control. β-actin was used as a reference gene since it is commonly used and none of the treatments were expected to affect its expression.

### 2.6 Phagocytosis Assay with Flow Cytometry

RAW 264.7 and Daudi cells were separately plated in 24 well plates at a density of 1 × 10^5^ cells/well each. The same day, designated RAW 264.7 wells were treated with PBI1 to reach a final concentration of 4.5 µg/mL in 0.5% DMSO. A slightly lower concentration was utilized in this experiment to demonstrate that profound changes in cell phenotype are possible at low concentrations of PBI1. 24 h following treatment, RAW cells were trypsinized using TrypLE (ThermoFisher), washed twice in PBS, and labelled with 10 µg/mL PE-F4/80 antibody (BD Biosciences, cat. No: 565410, Clone:T45-2342) for 30 min at 4 °C. All RAW cells were resuspended in culture media and 10 µM cytochalasin D was added to the designated samples. Simultaneously, Daudi cells were removed from the 24 well plate, washed twice with LCIS (ThermoFisher) and labelled with pHRhodo Green according to the manufacturer’s instructions (ThermoFisher). These cells were washed with PBS and then labelled with 10 µg/mL anti-CD47 antibody (Bio Xcell, cat. No: BE0270, Clone: MIAP301) for 30 min at 4 °C. All of the Daudi cells (approximately 2 × 10^5^) were resuspended in culture media and combined with the RAW 264.7 cells for 2 h at 37 °C. The approximate ratio of RAW:Daudi was 2:1. The samples were then washed with PBS, resuspended in FACS buffer (1% FBS in PBS) and transferred to flow cytometry tubes. The samples were analyzed on a LSRFortessa 5L flow cytometer (BD Biosciences) using 488 nm and 561 nm lasers, counting 10000 events, at the University of Massachusetts Amherst Flow Cytometry Facility. One sample per treatment group was evaluated. Phagocytotic index was calculated using the following equation: [F4/80^+^pHRhodoGreen^+^ events]/[F4/80^+^pHRhodoGreen^-^ events], normalized to the untreated control group.

### 2.7 Griess Assay

RAW 264.7 cells were plated in 24 well plates at a density of 1.5 × 10^5^ cells/well 24 h prior to the experiment. On the following day, the culture media was removed and replaced with serum free Opti-Mem media for 2 h. Cells designated for TLR4 inhibition were pre-treated with 7.2 µg/mL TAK242 (Cayman Chemical) 1 h before additional treatment. Then, media was replaced again with 250 µL phenol red-free DMEM culture media containing either 100 ng/µL LPS or 20 µL/mL PBI1 and cells were incubated for an additional 48 h. Cell supernatant was collected from wells and centrifuged at 5000 rpm for 5 min. Then, Griess reagent (Thermofisher) was prepared according to the manufacturer’s instructions and 60 µl Griess reagent was combined with 60 µL cell supernatant in a clear 96 well plate. After 15 min in the dark, absorbance was read using a SpectraMax M2 plate reader (Molecular Devices, Sunnyvale, CA) at 548 nm. Three biological replicates per treatment condition were used. Actual NO_2_^-^ concentrations were determined by comparing absorbance values to a standard curve generated using pure NaNO_2_^-^ solutions.

### 2.8 Confocal Microscopy

For acquisition of representative cell morphology images, RAW 264.7 cells were plated in a 4 chamber Lab-Tek II chambered cover-glass system (Nunc, NY) at a density of 5 × 10^4^/well and allowed to adhere overnight. Cells were then polarized with 20 µg/mL PBI1 in 0.2% DMSO for 48 h. After polarization, cells were imaged using an Eclipse Ti-E microscope at 20× magnification. M1 cell images were acquired similarly, using polarization conditions as outlined in the RT-PCR Preparation section above.

For acquisition of representative phagocytosis images, 5 × 10^4^ RAW 264.7 cells were plated in a 4 chamber Lab-Tek II chambered cover-glass system. Designated wells were treated with PBI1 to a final concentration of 4.5 µg/mL in 0.4% DMSO for 24 h. The RAW cells were labeled with Cell Tracker Blue according to the manufacturer’s instructions. 1 × 10^5^ Daudi cells were counted, labeled with phRhodo Green and added to the wells containing RAW cells. After 2 h incubation, wells were imaged using an Eclipse Ti-E microscope at 63× magnification. The final ratio of RAW to Daudi cells was approximately 2:1.

## 3. Results and Discussion

Following synthesis and characterization of PBI1 (**Fig. S1-S8**), RAW 264.7 murine macrophages were dosed with the compound, and changes in morphology and gene expression were assessed using confocal microscopy and RT-PCR, respectively (**Fig. 2**). Confocal microscopy revealed significant changes in macrophage morphology after 24 h. Treated macrophages acquired a phenotype that is associated with M1-polarized macrophages (**Fig. 2a**): the cells became flatter and produced longer pseudopodia [16] compared to non-treated cells. RT-PCR analysis of the expression of two M1-related markers, tumor necrosis factor-α (TNF-α) and inducible nitric oxide synthase (iNOS) [11], revealed that PBI1-treated macrophages express significantly higher levels of both genes compared to non-treated cells, in a similar trend as IFN-γ/LPS treatment (**Fig. 2b**). This effect was also observed in an immortalized primary macrophage cell line (**Fig. S9**). Furthermore, PBI1 treatment was also able to ‘re-educate’ macrophages that had been polarized toward the anti-inflammatory M2 phenotype, resulting in enhanced levels of iNOS and TNF-α. Alamar blue assays indicated that no significant toxicity occurred as a result of compound treatment (Fig. S10).

**Fig. 2.**
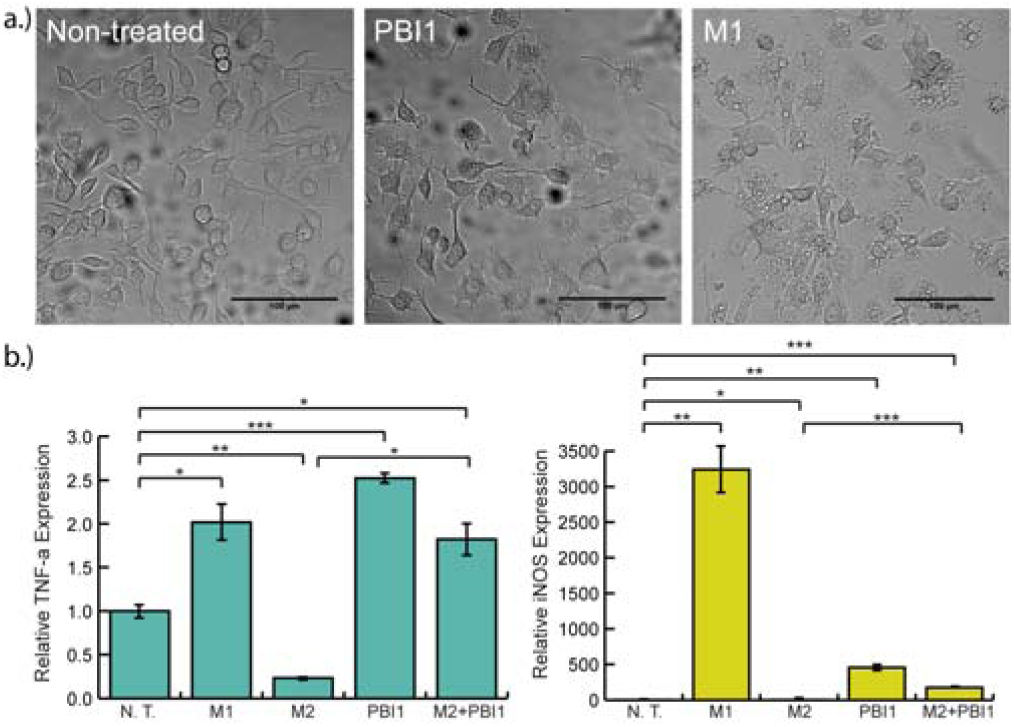
Effects of PBI1 treatment on cell morphology and M1 marker expression. a) Confocal microscopy images (20x) depicting changes in cell morphology following treatment with 20 µg/ml PBI1. PBI1-treated macrophages resembled those polarized to an M1 phenotype, with a “flatter” appearance and more pseudopodia, in comparison with non-treated cells. b) RT-PCR analysis of TNF-α and iNOS expression in PBI1 treated cells revealed significantly greater expression of both relative to non-treated controls. M2+PBI1 refers to cells polarized to the M2 phenotype and subsequently treated with PBI1. Unpaired, two-tailed student T-tests with equal variance were used to determine significance; *P ≤ 0.05, **P ≤ 0.005, ***P ≤ 0.0005.

Generation and release of reactive oxygen and nitrogen species (ROS and RNS, respectively) are important processes of M1 macrophage anti-tumor and pathogen responses [25]. RNS are generally derived from nitric oxide (NO). To evaluate relative NO production by PBI1-treated cells, a Griess assay was performed [26]. This method indirectly measures NO via evaluation of nitrite (NO_2_^-^), one of its two primary stable and non-volatile breakdown products. The determined NO_2_^-^ concentrations show that 20 µg/mL PBI1 effectively induced production of NO_2_^-^, achieving significantly higher levels than non-treated samples, and within 2-fold of (100 µg/mL) LPS-treated cells (**Fig. 3**). Additionally, pretreatment with the TLR4 specific inhibitor TAK242 completely eliminated the induction of NO by PBI1 (**Fig. 3**) [27]. This result demonstrates that PBI1 is an agonist of TLR4 and causes the activation of macrophages through this pathway.

**Fig. 3.**
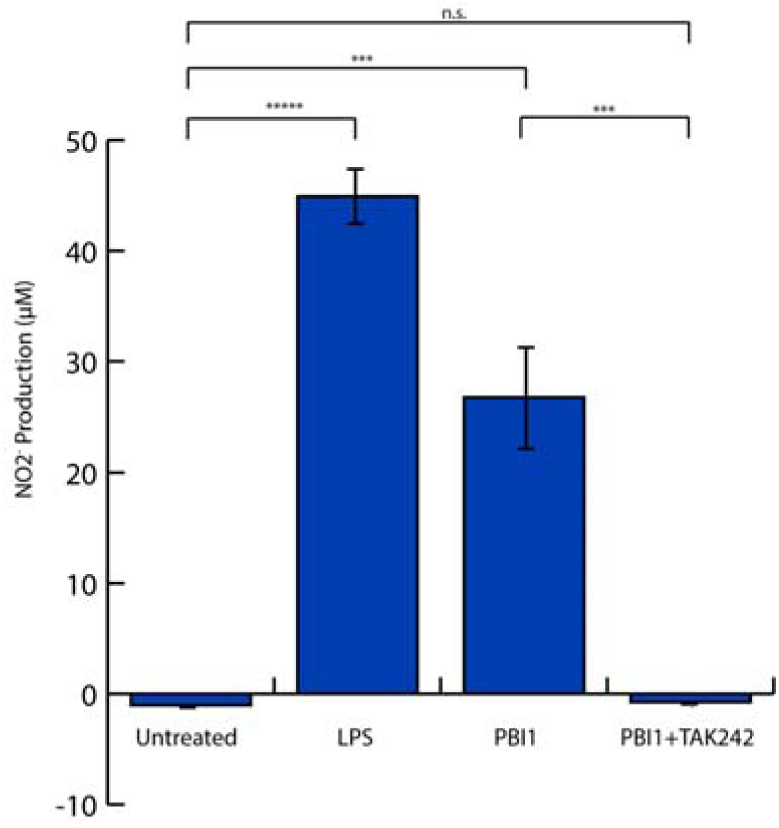
Supernatant NO_2_^-^ levels following LPS (100 µg/mL) or PBI1 (20 µg/mL) treatment as measured by Griess assay. Pretreatment with TAK242 inhibited TLR4 activity prior to PBI1 addition (PBI1+TAK242). Unpaired, two-tailed student T-tests with equal variance were used to determine significance; ***P ≤ 0.005, *****P ≤ 0.00005

A key aspect of the macrophage antitumor response is the identification and phagocytosis, or engulfment, of cancerous cells [6]. To assess the phagocytic efficacy of PBI1 treated macrophage cells, a fluorescent flow cytometry assay was conducted using Daudi (B cell lymphoma) as the target cancer cell line (**Fig. 4**, **S11**). RAW 264.7 cells treated with PBI1 had a nearly 5-fold greater phagocytic efficiency versus non-treated macrophages. This was increased further when macrophage PBI1 treatment was combined with antibody-mediated blocking of Daudi cell CD47, the ‘don’t eat me’ signal involved in phagocytosis inhibition signaling. Cytochalasin D, a potent phagocytosis inhibitor, was used as a negative control [28].

**Fig. 4.**
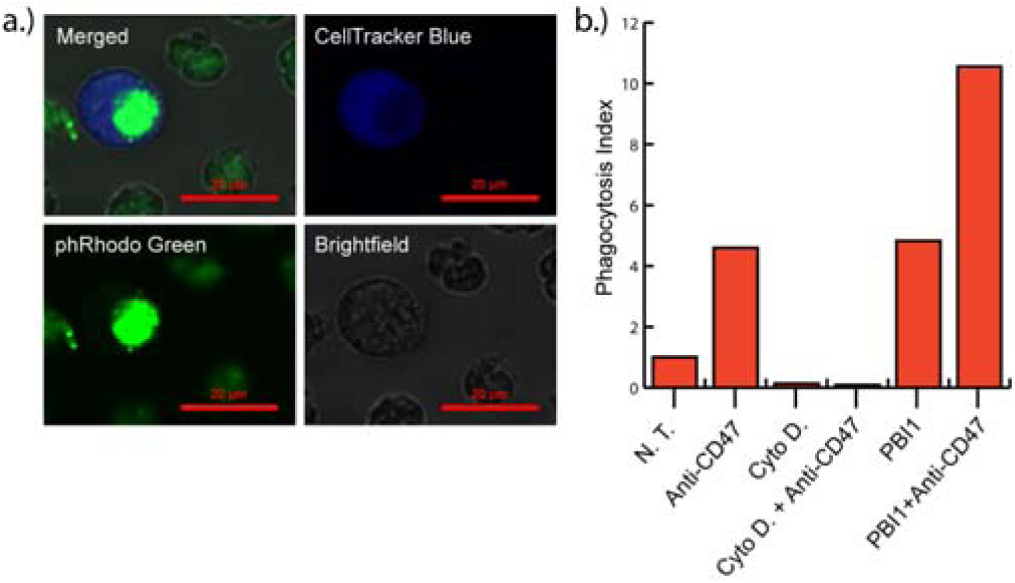
Effects of PBI1 on phagocytosis of Daudi (B-cell lymphoma) cells. a) Representative confocal microscopy images illustrating RAW 264.7 macrophage (CellTracker Blue) phagocytosis of Daudi cells (phRodo Green). b) Phagocytosis indices generated from flow cytometry data of RAW 264.7 macrophages phagocytosing Daudi cells following various treatments.

In summary, PBI1 was demonstrated to polarize macrophages toward an anti-cancer phenotype. RAW 264.7 macrophages treated with PBI1 adopted an “M1-like” pro-inflammatory morphology, as cells became flattened and produced extended pseudopodia. RT-PCR results also corroborated this phenotype via an increase in the expression of M1 inflammatory genes. PBI1 treatment increased the expression of TNF-α and iNOS approximately 2.5 and 500-fold, respectively. Dosing of PBI1 also resulted in the re-education of M2-polarized macrophages, whereupon following treatment, cells expressed higher levels of inflammatory cytokines. This is particularly relevant for the conversion of tumor-associated macrophages into tumor-killing macrophages within cancer microenvironments.

We also evaluated the mechanism by which PBI1 promoted macrophage activation. The most common pathways for macrophage stimulation include the TLR family of receptors that recognize a wide variety of substrates [8]. To confirm that TLR4 is the target of PBI1 for activation, a competition experiment was performed. Cells were treated with a highly specific TLR4 inhibitor, TAK242, and activation by PBI1 was determined by Griess assay. This experiment revealed that macrophages treated with either PBI1 or LPS released significant levels of nitric oxide, another confirmation of the activation potential of PBI1. However, with pre-treatment of cells with TAK242, PBI1 did not induce any detectable level of nitric oxide, confirming that TLR4 is the target of PBI1 and is crucial to resulting macrophage activation.

Having evaluated the pro-inflammatory responses of macrophages to PBI1 treatment, the ability of these cells to subsequently phagocytose cancers cells was investigated. It has been previously shown that activation of local macrophages into an M1 inflammatory phenotype can result in significant anti-tumor macrophage activity, which is of great interest for the generation of cancer immunotherapies [24]. One of the major anti-cancer macrophage mechanisms involves phagocytosis of cancer cells. Inflammatory macrophages invade can tumor tissue, engulf resident tumor cells, and release immune signaling factors that drive further immune responses against the tumor site [24]. A phagocytosis assay revealed that PBI1-treated macrophages engulfed targeted cancer cells with a 5-fold higher efficacy than untreated macrophages, revealing that inflammatory activation by PBI1 does in fact increase anti-tumor activity. It was also demonstrated that pre-treatment of the targeted cancer cells with a CD47 blocking antibody, which blocks the SIRPα-CD47 phagocytosis inhibitory pathway, further increases the efficacy of PBI1-induced phagocytosis. This effect occurred independently of PBI1-activation as well and could be used to increase the anti-cancer efficacy of PBI1 treatment.

## 4. Conclusion

The immune system and cancer progression have a complex relationship. While effective immune-based strategies are of interest in enervating and eliminating cancer progression, their development can be challenging due to varying macrophage-activating signals [29,30]. PBI1 has significant macrophage activation capability. As a potential immune adjuvant for cancer therapy, PBI1 is relatively non-toxic and effectively increases both phagocytic and oxidative burst mechanisms of the macrophage anti-tumor response. While PBI1 has shown promise as a therapeutic on its own, it will likely show the greatest efficacy if used in conjugation with drug delivery vehicles or targeting elements. This is especially relevant in trying to avoid broad immune-activation and runaway immune responses. As a small molecule, it should be fairly straight-forward to attach to a variety of carriers, including nanoparticles, proteins, and cells; further studies will explore these options. Incorporation of PBI1 with more complex therapeutics, as opposed to its use independently, could result in robust combinatorial anti-cancer strategies. Additional studies will evaluate the anti-cancer efficacy of PBI1 *in vivo*, including in concert with delivery vehicles, and as an adjuvant with other chemotherapeutics. In the future, we will also seek to identify other potential therapeutic targets of PBI1-activated macrophages.

## ABBREVIATIONS

CD47: cluster of differentiation 47
IL-4: interleukin 4
CS1: CCND3 subset 1
CSF1R: colony stimulating factor receptor 1
MFEG8: Milk fat globule-EGF factor 8 protein
VEGF: Vascular endothelial growth factor
SEAP: Secreted embryonic alkaline phosphatase
LPS: Lipopolysaccharide
IFN-γ: Interferon-γ

## Author Contributions

J.M.H. and J.A.M. designed the study with input from M.E.F. S.H. synthesized and chemically characterized PBI1. J.M.H., J.A.M. and E.R. performed experiments and performed data analysis in conjunction with M.E.F. J.M.H. wrote and M.E.F. revised the manuscript. All authors reviewed and commented on the manuscript.

## ACKNOWLEDGMENT

This research was supported by a SEED grant from the University of Massachusetts Institute for Applied Life Sciences. JH was supported by the NIH (EB014277) and an NIH National Research Service Award (T32 GM008515). JAM-R was supported by the UMass Amherst NIH Postbaccalaureate Research Program (GM086264-08) and a Northeast Alliance for Graduate Education and the Professoriate (NEAGEP) fellowship from the STEM Diversity Institute at UMass Amherst.

## ABBREVIATIONS

CD47: cluster of differentiation 47
IL-4: interleukin 4
CS1: CCND3 subset 1
CSF1R: colony stimulating factor receptor 1
MFEG8: Milk fat globule-EGF factor 8 protein Milk fat globule-EGF factor 8 protein
VEGF: Vascular endothelial growth facto Vascular endothelial growth factor
SEAP: Secreted embryonic al-kaline phosphatase Secreted embryonic alkaline phosphatase

## Entry for the Table of Contents

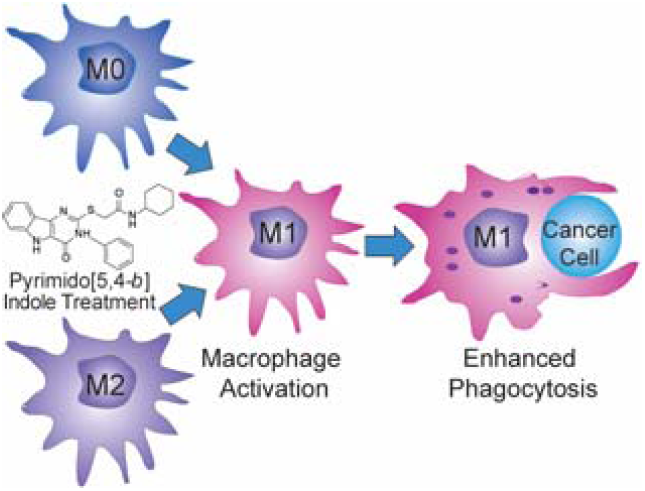
PBI1 can polarize undifferentiated M0 macrophages or re-program tumor-promoting M2 macrophages into the anti-tumor (M1) phenotype and enhance the phagocytosis of cancer cells.

