## Supplementary Information for "Macrophage Activation by a Substituted Pyrimido[5,4-*b*]Indole Increases Anti-Cancer Activity"

### Cell Viability/PBI1 Toxicity Assay

RAW 264.7 cells were plated in 24 well plates at a density of 5 x $10^4$ cells/well. PBI1 was dissolved in DMSO at varying concentrations and added to the wells at a final DMSO concentration of 0.4%. 24 h later, Alamar Blue Assay was performed according to the manufacturer's instructions and the fluorescence was determined using a SpectraMax M2 plate reader.

### Synthesis of PBI1

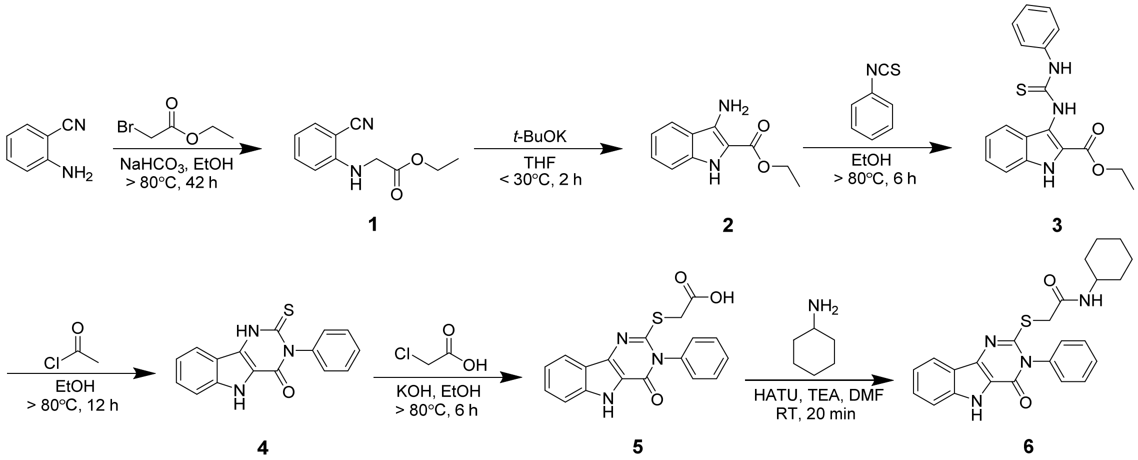


**Fig. S1 PBI1 Synthesis Scheme**

#### Ethyl 2-((2-Cyanophenyl)amino)acetate (1)

2-Aminobenzonitrile (11.8 g, 100 mmol), ethyl bromoacetate (11.5 mL, 104 mmol), and sodium bicarbonate (10 g, 117 mmol) were dissolved in anhydrous EtOH (30 mL) and refluxed for 48 h. After allowing to cool briefly, the precipitate was filtered, and following further cooling to rt, crystals were filtered and then recrystallized using ethanol. Crystals were washed with cold water to obtain a white crystalline product to give 12 g compound **1** in 70 % yield. **$^1$H NMR (400 MHz, DMSO-d6)** δ ppm 1.18−1.22 (m, 3 H), 4.05 (d, J = 6.41 Hz, 2 H), 4.13 (q, J = 7.22 Hz, 2 H), 6.34 (t, J = 6.25 Hz, 1 H), 6.65 (d, J = 8.54 Hz, 1 H), 6.70 (t, J = 7.47 Hz, 1 H), 7.41 (t, J = 7.78 Hz, 1 H), 7.49 (dd, J =7.93, 1.53 Hz, 1 H).

#### Ethyl 3-Amino-1H-indole-2-carboxylate (2)

In a flame-dried flask, a suspension of potassium t-butoxide (4.48 g, 40 mmol) in anhydrous THF was stirred and maintained below 30 °C under nitrogen atmosphere.

To this solution was added a solution of compound **1** (10 g, 40 mmol) in anhydrous THF over 45 min, which was then stirred for an additional 2 h. Cold water was added to the reaction, extraction was performed with EtOAc, and product dried over $MgSO_4$. The solids were then dissolved in minimal EtOH, and water was added to form a precipitate at room temperature. The precipitate was filtered and washed with cold EtOH to give 7.46 g compound **2** in 74.6 % yield. **$^1$H NMR (400 MHz, DMSO-d6)** δ ppm 1.34 (t, J = 7.02 Hz, 3 H), 4.30 (q, J = 7.12 Hz, 2 H), 5.68 (s, 2 H), 6.88 (ddd, J = 8.08, 5.80, 1.98 Hz, 1 H), 7.20−7.22 (m, 2 H), 7.74 (d, J = 7.93 Hz, 1 H), 10.34 (s, 1 H).

**Ethyl 3-(3-Phenylthioureido)-1H-indole-2-carboxylate (3)**
To a stirring solution of compound **2** (7 g, 33.6 mmol) in warm EtOH (50 mL), phenyl isothiocyanate (4.5 mL, 37.7 mmol) dissolved separately in warm EtOH was added dropwise. The reaction refluxed for 6 h, and cooled to rt overnight. The desired product precipitated, was filtered and washed with EtOH, and dried overnight *in vacuo* to give 9 g compound **3** in 77 % yield. **$^1$H NMR (400 MHz, DMSO-d6)** δ ppm 1.34 (t, J = 7.07 Hz, 3 H), 4.33 (q, J = 7.31 Hz, 2 H), 7.11 (dt, J = 14.75, 7.50 Hz, 2 H), 7.30−7.32 (m, 3 H), 7.45 (d, J = 8.29 Hz, 1 H), 7.52 (d, J = 7.80 Hz, 2 H), 7.58 (d, J = 8.29 Hz, 1 H), 9.41 (s, 1 H), 9.71 (br s, 1 H), 11.80 (s, 1 H).

**3-Phenyl-2-thioxo-2,3-dihydro-1H-pyrimido[5,4-b]indol-4(5H)-one (4)**
In a dried flask, acetyl chloride (7.45 mL, 106 mmol) was dissolved in 30 mL cold, anhydrous EtOH. Separately, compound **3** (7 g, 20.5 mmol) was dissolved in 75 mL anhydrous EtOH in a flame-dried flask charged with nitrogen, to which the acetyl chloride solution was then added. The reaction was refluxed for 12 h and then allowed to cool to rt. The precipitate was filtered and recrystallized with EtOH to give 5.25 g compound **4** in 86 % yield. **$^1$H NMR (400 MHz, DMSO-d6)** δ ppm 7.19−7.29 (m, 3 H), 7.29−7.52 (m, 6 H), 8.24 (d, J = 8.07 Hz, 1 H), 12.19 (br s, 1 H). 294.0698.

**2-((4-Oxo-3-phenyl-4,5-dihydro-3H-pyrimido[5,4-b]indol-2-yl)-thio)acetic Acid (5)**
Compound **4** (5 g, 17.9 mmol) and KOH (2 g, 35.86 mmol) were dissolved in 10 mL warm anhydrous EtOH in a flame-dried flask. Separately, chloroacetic acid (1.74 g, 17.93 mmol) was dissolved in 100 mL anhydrous EtOH in a flame-dried flask. The chloroacetic acid solution was then added to the compound **4-**containing mixture, and refluxed for 6 h. After concentrating the reaction by half *in vacuo*, the solution was acidified to pH 4 using 3M HCl. The resulting precipitate was filtered and rinsed with water, to give 4.6 g compound **5** in 78 % yield. **$^1$H NMR (400 MHz, DMSO-d6)** δ ppm 3.98 (s, 2 H), 7.24− 7.28 (m, 1 H), 7.46−7.63 (m, 7 H), 7.9 (d, J = 7.70 Hz, 1 H), 12.10 (br s, 1 H).

**N-Cyclohexyl-2-((4-oxo-3-phenyl-4,5-dihydro-3H-pyrimido[5,4-b]indol-2-yl)thio)acetam ide (6; PBI1)**
Compound **5** (1 mg, 2.8 mmol), triethylamine (800 µL, 5.6 mmol), and cyclohexylamine (400 µL, 3.2 mmol) were dissolved in anhydrous DMF (20 mL). To

this mixture, HATU (1.2 g, 3.2 mmol) dissolved in 1 mL of DMF was added. The reaction was stirred for 20 min at ambient temperature and then concentrated *in vacuo*. The crude material was recrystallized with MeOH to give 800 mg in 65% yield. **$^1$H NMR (400 MHz, DMSO-d6)** δ ppm 1.19-1.21 (m, 4 H), 1.52 (d, J = 8.53 Hz, 1H ,), 1.63−1.73 (m, 4 H), 3.51 (m, 1 H, *H*-6), 3.89 (s, 2 H), 7.25 (t, J = 7.29 Hz, 1 H), 7.45−7.61 (m, 6 H,), 8.06 (d, J = 7.98 Hz, 1 H), 8.16 (d, J = 7.70 Hz, 1 H), 12.09 (s, 1 H). FTMS m/z (ESI) calc. for $C_{24}H_{24}N_4O_2Sna$ $(M + Na)^+$, 455.1512; found, 455.1511.

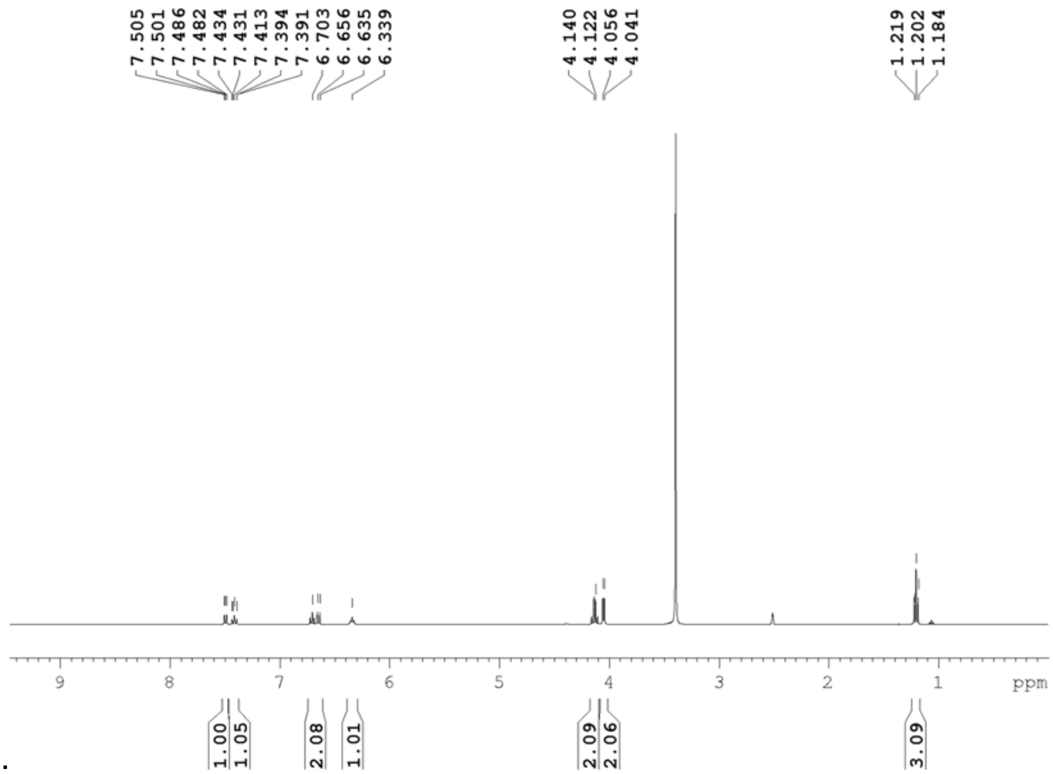


**Fig. S2 $^{1}$H NMR spectrum of Ethyl 2-((2-Cyanophenyl)amino)acetate (1)**

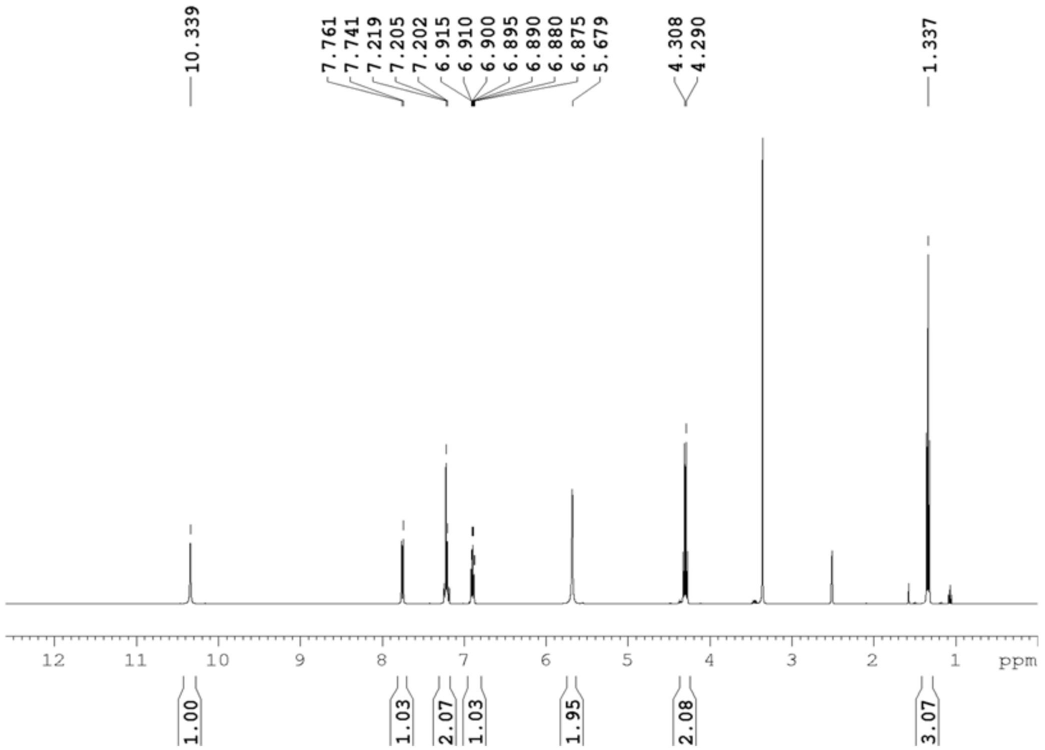


**Fig. S3 $^{1}$H NMR spectrum of Ethyl 3-Amino-1H-indole-2-carboxylate (2)**

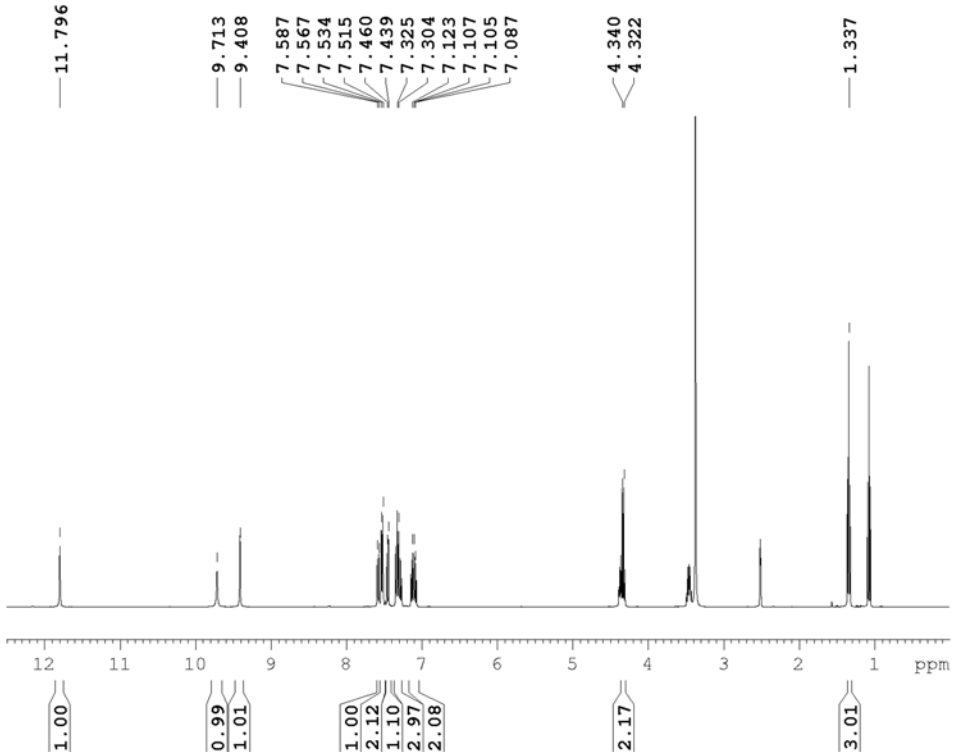


**Fig. S4. ¹H NMR spectrum of Ethyl 3-(3-Phenylthioureido)-1H-indole-2-carboxylate (3)**

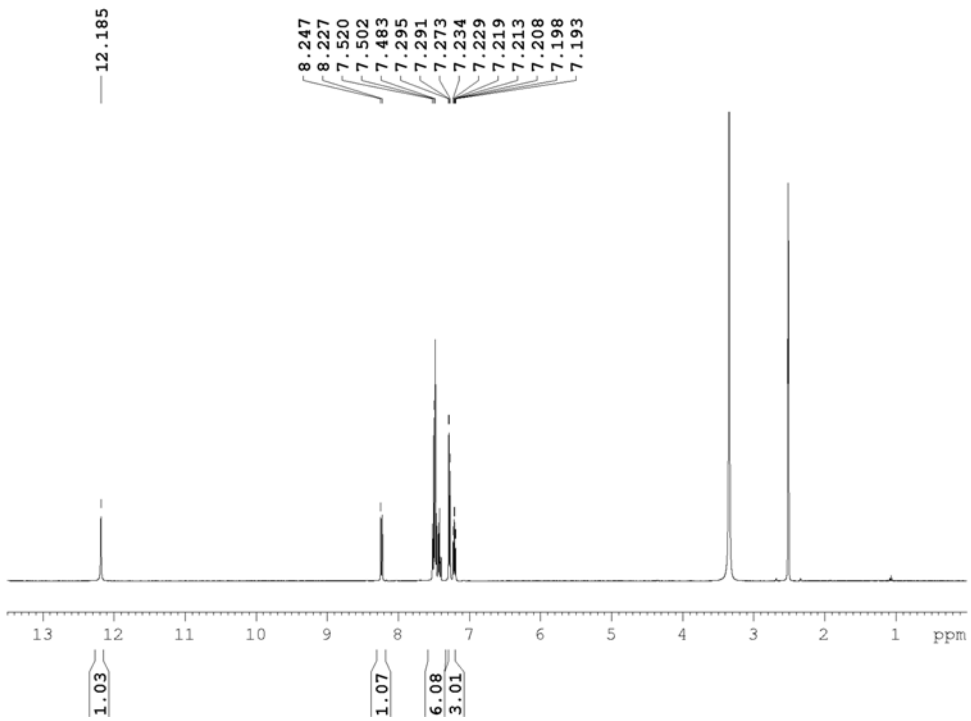


**Fig. S5 ¹H NMR spectrum of 3-Phenyl-2-thioxo-2,3-dihydro-1H-pyrimido[5,4-b]indol-4(5H)-one (4)**

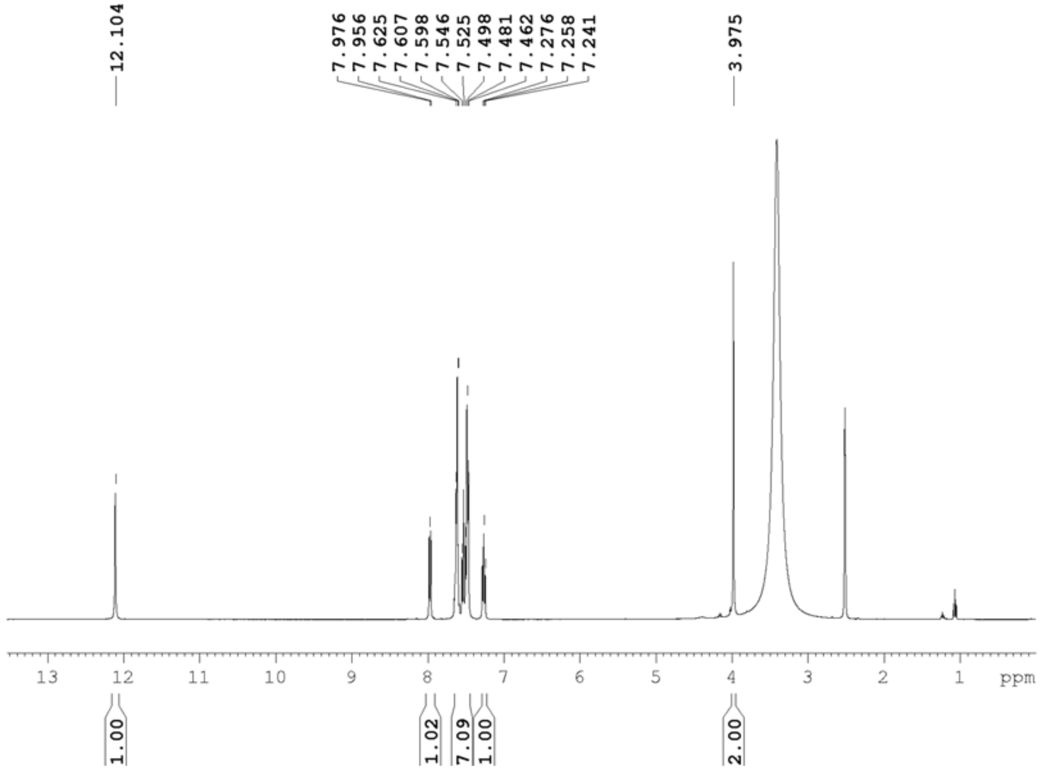


**Fig. S6 $^1$H NMR spectrum of 2-((4-Oxo-3-phenyl-4,5-dihydro-3H-pyrimido[5,4-b]indol-2-yl)-thio)acetic Acid (5)**

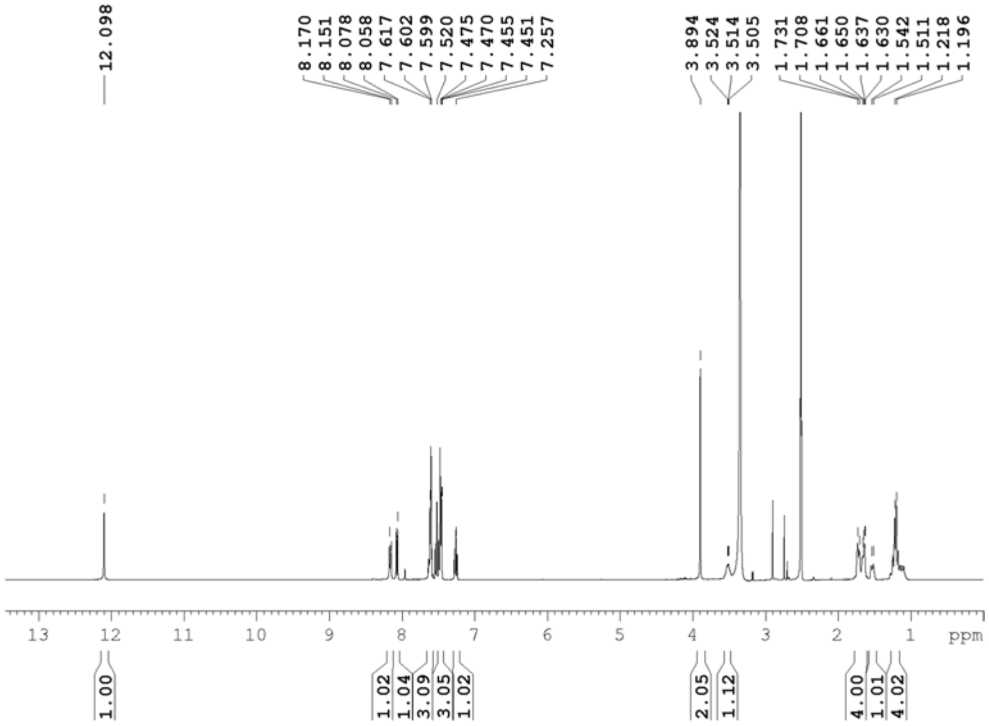


**Fig. S7 $^1$H NMR spectrum of N-Cyclohexyl-2-((4-oxo-3-phenyl-4,5-dihydro-3H-pyrimido[5,4-b]indol-2-yl)thio)acetamide (6)**

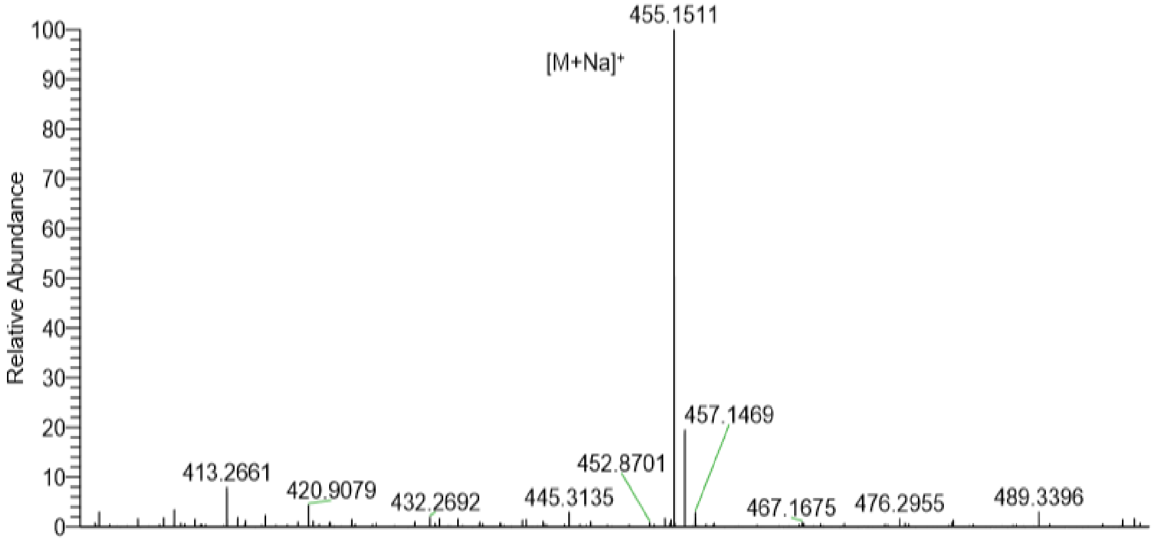


**Fig. S8 FTMS spectrum of N-Cyclohexyl-2-((4-oxo-3-phenyl-4,5-dihydro-3H-pyrimido[5,4-b]indol-2-yl)thio)acetamide (6)**

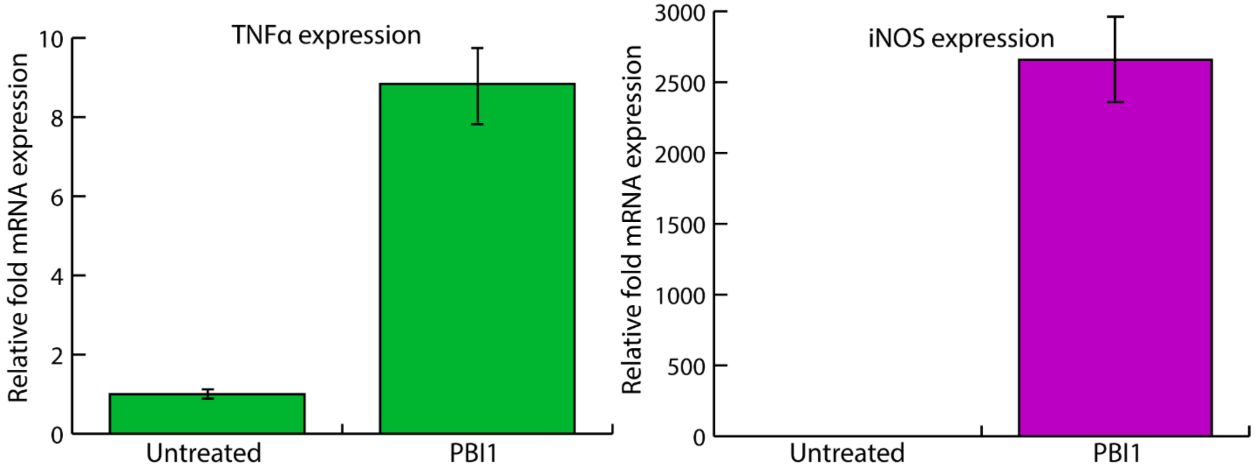


**Fig. S9 RTPCR analysis of M1 related genes in primary, immortalized macrophages (PIMs) following PBI1 treatment** PIMs treated with PBI1 display increased levels of TNFα and iNOS expression as analyzed by RT-PCR. Three biological replicates with three technical replicates each were used in analyzing each treatment condition. Error bars represent standard deviation.

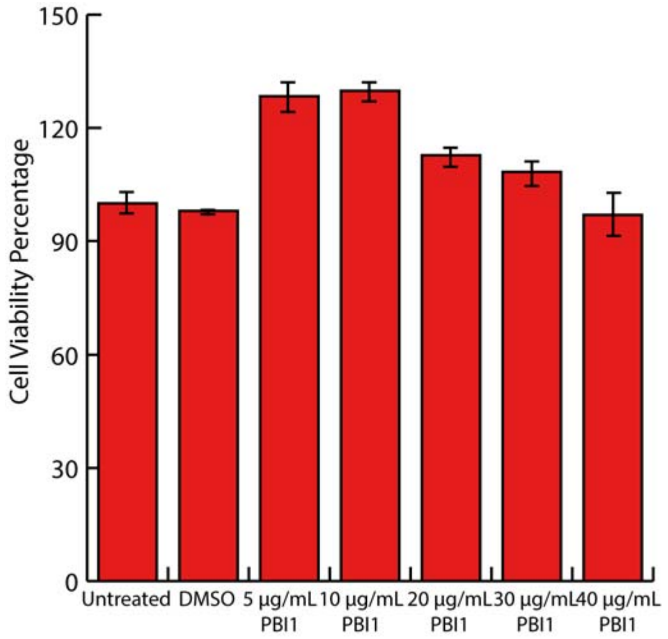


**Fig. S10 Toxicity of PBI1 as evaluated by Alamar Blue Assay** RAW 264.7 macrophages were treated with increasing amounts of PBI; subsequent cellular viability was assessed using Alamar blue reagent. Minimal differences in viability were observed at the utilized concentrations. Three biological replicates were included per treatment condition; error bars represent standard deviation.

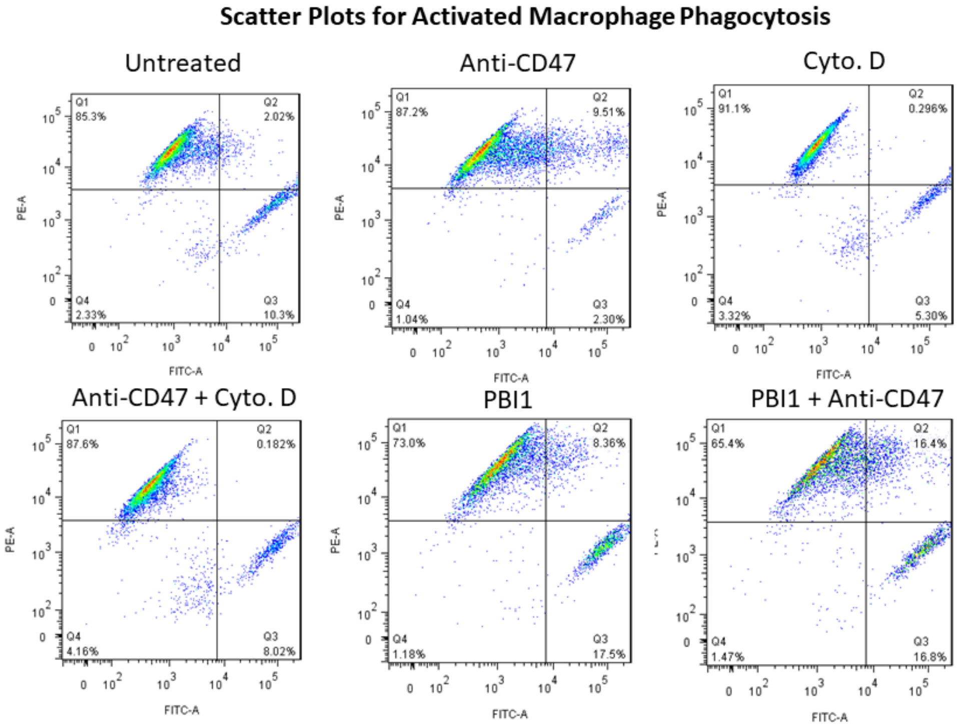


**Fig. S11 Representative flow cytometry scatter plots for analysis of macrophage phagocytosis** For analysis of these data, the quadrant gate was placed relative to populations in the "untreated" group. The vertical axis was positioned at the right-most extreme of the PE stained macrophage population, and the horizontal axis was positioned on the top extreme of the FITC positive Daudi population. Q2 was identified as $PE^{+}FITC^{+}$ events (macrophage phagocytosing cancer cells). Each scatter plot includes 10,000 events.
